# circMRPS35 promotes malignant progression and cisplatin resistance in hepatocellular cancer

**DOI:** 10.1101/2021.03.24.436699

**Authors:** Peng Li, Runjie Song, Huijiao Liu, Mei Liu, Fan Yin, Shuoqian Ma, Xiaomeng Jia, Xiaohui Lu, Yuting Zhong, Xiru Li, Xiangdong Li

## Abstract

Hepatocellular carcinoma (HCC), a common malignant tumor, is one of the main causes of cancer-related deaths worldwide. Circular RNAs (circRNAs), a novel class of non-coding RNA, have been reported to be involved in the etiology of various malignancy. However, the functions of circRNAs in HCC remain unclear. In this study, through mining the RNA sequencing databases from GEO datasets and subsequent experimental verification, we identified that hsa_circ_0000384 (circMRPS35) was highly expressed in HCC. Knockdown of circMRPS35 suppressed the proliferation, migration, invasion, clone formation and cell cycle of HCC cell lines both in vitro and in a xenograft mouse model. Mechanically, circMRPS35 sponged microRNA-148a-3p (miR-148a), which in turn regulated STX3-PTEN axis. Surprisingly, we detected a peptide encoded by circMRPS35 (circMRPS35-168aa), which was significantly induced by chemotherapeutic drugs and promoted cisplatin resistance in HCC cells. These results demonstrated that circMRPS35 might be a novel factor in HCC progress, and has a great potential as a new diagnosis and therapeutic target for treatment of HCC.

## Introduction

Hepatocellular carcinoma (HCC) is one of the most frequently diagnosed cancers and cancer-related deaths globally ^1–3^. Due to the lack of symptoms in the early stage of HCC, most patients are usually diagnosed at advanced stage, and the 5-year survival rate is approximately 14% for HCC patients ^4, 5^. Therefore, the valuable diagnostic biomarkers and therapeutic targets are urgently needed to be explored and verified. In general, surgical resection combined with chemotherapy is curative for the early stage of HCC ^6^. However, chemoresistance was detected in most HCC patients with long-term chemotherapy, leading to the poor prognosis ^7, 8^. Therefore, the molecular mechanism of chemoresistance in HCC is needed to be further studied.

Circular RNAs (circRNAs) serve as one types of non-coding RNAs which are covalently closed signal-stranded RNAs derived from the back-spliced mechanism of pre-mRNA during the process of transcription ^9, 10^. Recently, with the advance of sequencing technologies and bioinformatics approaches, more and more circRNAs were found and some of them were proved with the significant bio-functions ^11^. A number of circRNAs play important biological roles in HCC process ^12–14^. Studies have showed that the unusually expressed circRNAs influenced the tumorigenesis with multiple functions.

In this study, by re-analyzing the RNA sequencing database from GEO datasets (GSE77509, GSE114564 and GSE159220) combined with experimental verification, we observed that hsa_circ_0000384 (circMRPS35) was significantly elevated in HCC. We hypothesized that circMRPS35 might have a crucial role in HCC progression. To test our hypothesis, we used stable circMRPS35 silenced Huh-7 and HCC-LM3 cell lines to address its critical roles in cell growth and invasion during tumorigenesis both in vitro and in vivo. Surprisingly, we also found that circMRPS35 encoded a novel peptide with 168 amino acids induced by chemotherapeutic drugs, which promoted HCC cells resistance to cisplatin treatment. Our findings may provide a better understanding of the clinical significance of circMRPS35, which implied that circMRPS35 might be a new diagnosis and therapeutic target for the treatment of HCC.

## Result

### The expression and characteristics of circMRPS35 in HCC tissues and cell lines

To find the differentially expressed circRNAs between HCC and adjacent samples, we mined the RNA sequencing database from three GEO datasets (GSE77509, GSE114564 and GSE159220). After re-analysis, we selected 8 markedly differential expressed circRNAs in all the three datasets (Fig.1A, S1A and Table S2-4). Due to 4 of these 8 circRNAs were deeply reported in HCC ^15–18^, we then detected the expressions of the other 4 circRNAs by using 10 pairs of human tissues (HCC tissues vs. the correspondent non-tumor adjacent tissues). We found that circMRPS35 was the most significantly different expressed in HCC tissues (Fig.S1B). By comparing the expressions of HCC cell lines (HepG2, SMMC-7721, Huh-7, HCC-LM3, SNU-398) and the normal liver cell line (L02), together with the other 25 pairs of human HCC samples, we further confirmed that circMRPS35 was highly up-regulated in both HCC cell lines and the HCC tissues (Fig. 1B and C). Furthermore, we performed receiver operating characteristic (ROC) analysis to evaluate the diagnostic value of circMRPS35, and the result showed that the sensitivity of diagnosis was high (value of the area under the ROC curve (AUC) was 0.8147) in HCC (Fig. 1 D).

**Figure 1.**
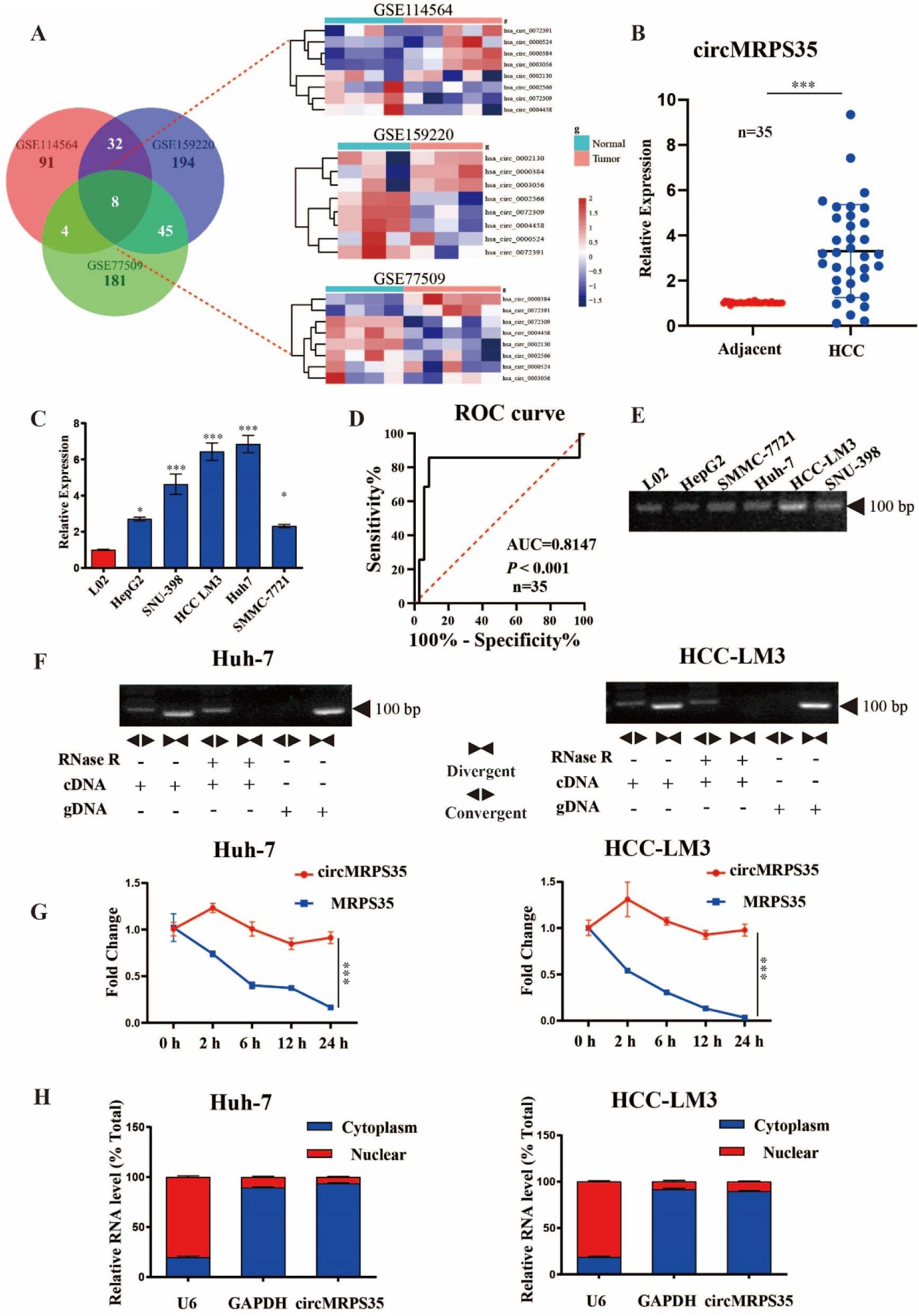
The expression and characteristics of circMRPS35 in HCC tissues and cells. (A) Schematic illustration showing the significantly different expressions circRNAs predicted by overlapping GSE77509, GSE114564 and GSE159220 data (left) and expression heat map of those overlapping circRNAs (right). (B) RT-qPCR analysis of circMRPS35 in 35 pairs of HCC and adjacent tissues. (C) RT-qPCR analysis of circMRPS35 in HCC cell lines compared to L02 cells. (D) The ROC curve of the diagnostic value of circMRPS35. (E) RT-PCR analysis of circMRPS35 in HCC cell lines and L02 cells. (F) RT-PCR analysis of circMRPS35 and MRPS35 with divergent and convergent primers after RNase R treatment. (G) RT-qPCR analysis of circMRPS35 and MRPS35 after ACTD treatment. (H) RT-qPCR analysis of circMRPS35 after RNA Nucleocytoplasmic separation, U6 and GAPDH as markers of nucleus and cytoplasm, respectively. Error bars represent the means ± SEM of 3 independent experiments. \**P* < 0.05, \*\**P* < 0.01, \*\*\**P* < 0.001.

After the bioinformatic analysis in the circBase database, we observed that circMRPS35 was derived from a mitochondrial ribosomal protein S35 *(MRPS35)* with exon 2 to exon 5 (410 bp) of head-to-tail back-spliced (Fig. S1C). By using a pair of divergent primers crossing the splicing site, we found a band (130 bp) of circMRPS35 in HCC cells and L02 cells by reverse transcription PCR (RT-PCR) (Fig. 1E). Furthermore, we observed that RNase R enzyme treatment could not destroy the cyclic structure of circMRPS35, compared with the liner transcription of *MRPS35* in HCC-LM3 and Huh-7 cells (Fig. 1F). Moreover, we noticed that circMRPS35 had a longer half-life than the linear transcript of MRPS35 in both HCC-LM3 and Huh-7 cells upon actinomycin D (ACTD) treatment (Fig. 1G). Next, the back-spliced sites of circMRPS35 were confirmed by Sanger sequencing (Fig. S1D). By using nucleus-cytoplasmic separation analysis, we found that circMRPS35 was predominantly localized in the cytoplasm of HCC-LM3 and Huh-7 cells (Fig. 1H), respectively.

These results suggested that circMRPS35 was highly expressed in HCC and predominantly located in the cytoplasm of HCC cells.

### CircMRPS35 acts as an oncogene in HCC cells

To further study the molecular actions of circMRPS35 in HCC cells, we silenced the expression of circMRPS35 by the short hairpin RNAs (shRNAs) against the back-spliced sites of circMRPS35 (Fig. 2A). By using the lentivirus system, we found that circMRPS35 was knocked down significantly, meanwhile we also confirmed that these circMRPS35 specific shRNAs did not affect the liner transcription of *MRPS35* in Huh-7 and HCC-LM3 cells by real-time quantitative PCR (RT-qPCR) (Fig. 2B), respectively. Next, by using the cell viability and colony formation assays, we demonstrated that the proliferations of Huh-7 and HCC-LM3 cells were suppressed significantly when circMRPS35 was stably silenced (Fig. 2C and D). Subsequently, the wound healing and transwell assays showed that the cell migration and invasion of the stable circMRPS35 silenced Huh-7 and HCC-LM3 cells were significantly slowed down, compared to the Huh-7 or HCC-LM3 control cells (Fig. 2E and F), respectively. For cell cycle progression, the results of flow cytometry showed a significant increase in the number of cells in the G0/G1 phase and a concomitant reduction in G2/M phase in the stable circMRPS35 silenced Huh-7 and HCC-LM3 cells (Fig. 2G and H). To evaluate the biological functions of circMRPS35 in vivo, stable circMRPS35 silenced or corresponding control Huh-7 and HCC-LM3 cells were subcutaneously injected into the BALB/c nude mice, respectively (n = 6). The growth rate and size (volume and weight) of the xenograft tumors in stable silenced circMRPS35 groups were decreased compared to the Huh-7 and HCC-LM3 control groups (Fig. 2I), respectively. Immunohistochemistry (IHC) analysis of the xenograft tumors tissues showed that Ki67 was highly expressed in the control tumors, compared to the stable circMRPS35 silenced tumors (Fig. 2J).

**Figure 2.**
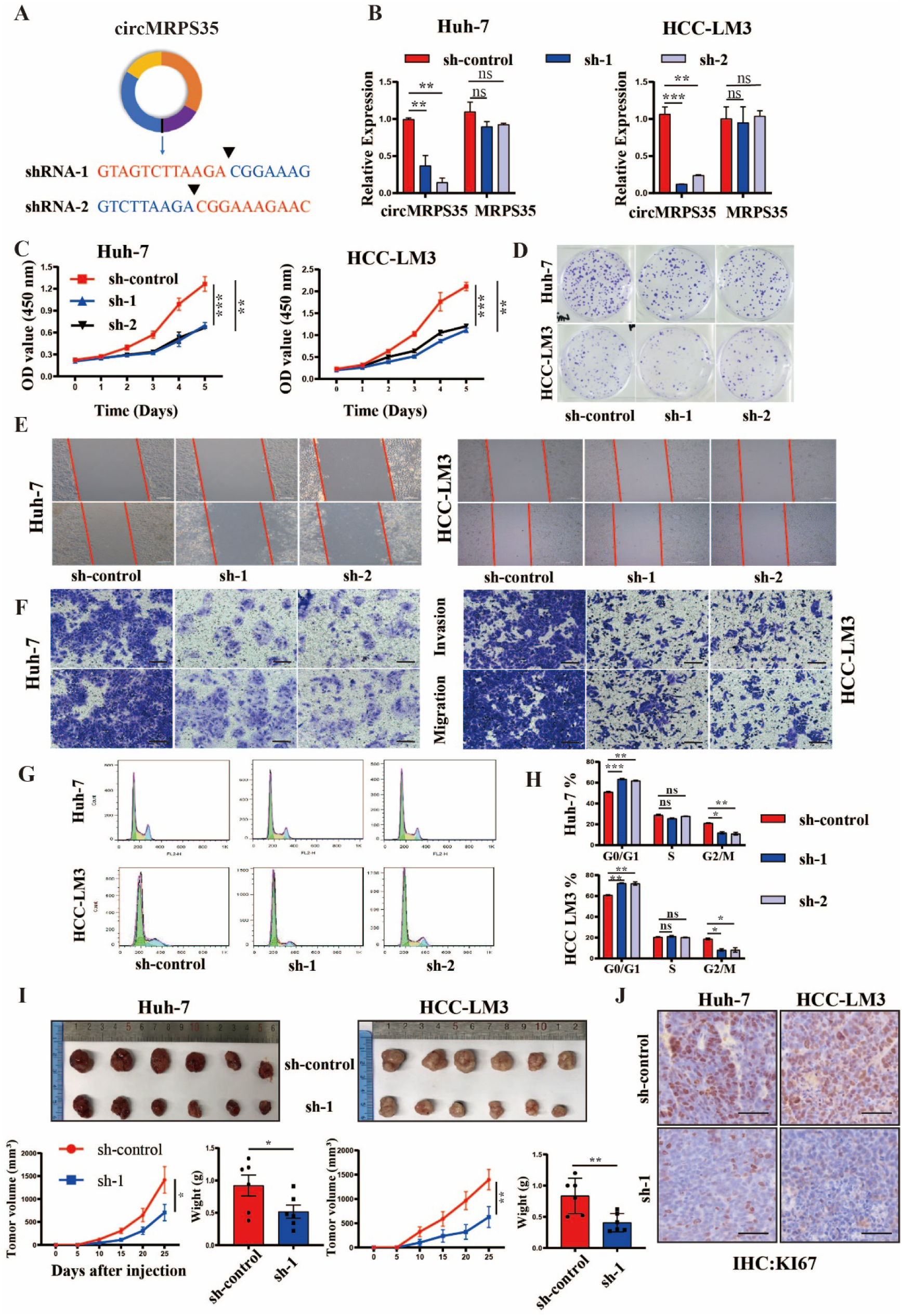
CircMRPS35 acts as an oncogene in HCC cells. (A) Schematic representation of target sequences about shRNAs of circMRPS35. (B) RT-qPCR analysis of circMRPS35 and MRPS35 of circMRPS35 silenced HCC-LM3 and Huh-7 cells. (C) Cell viability assays were used to test proliferation of HCC-LM3 and Huh-7 after silencing of circMRPS35. (D) Colony formation assays were performed to test cell growth of HCC-LM3 and Huh-7 cells after silencing of circMRPS35. (E) Wound healing experiments were used to detect cell migration of LM3 and Huh-7 cells after silencing of circMRPS35. (F) Transwell assays of invasion and migration in circMRPS35 silenced HCC-LM3 and Huh-7 cells. (G-H) Cell cycle assays were used to detect cell cycle arresting level of circMRPS35 silenced HCC-LM3 and Huh-7 cells. (I) BALB/c nude mice (n = 6 each group) were injected circMRPS35 silenced or control HCC-LM3 and Huh-7 cells. Sizes of xenografted tumors were measured every 5 days and weights of xenografted tumors were summarized after being sacrificed. (J) IHC analysis of Ki67 for circMRPS35 silenced or control HCC-LM3 and Huh-7 xenograft tumors tissues. Error bars represent the means ± SEM of 3 independent experiments. \**P* < 0.05, \*\**P* < 0.01, \*\*\**P* < 0.001.

Taken together, the results showed that low expression of circMRPS35 inhibited the proliferation, migration, invasion, cell cycle of HCC cells, and tumor growth both in vitro and the xenograft tumor models in vivo.

### CircMRPS35 serves as a sponge for miR-148a in HCC cells

CircRNAs can serve as microRNA’s sponge through the complementary binding sites ^19^. As circMRPS35 is located in the cytoplasm of HCC cells, we explored whether circMRPS35 promoted HCC progress through interacting with microRNA (miRNA). To predict and screen the possible miRNA candidates, we assessed multiple bioinformatics programs (miRanda, ENCORI and circBank) and selected a list of 24 potential miRNAs that might bind to circMRPS35 directly (Fig. 3A and Table S5). In addition, by using the cancer genome atlas (TCGA) database, we screened out the expression patterns of the selected miRNAs in HCC patients (Fig. S2A). Of particular, based on the results from the expressions and the prognosis of this list of miRNAs, we selected 4 highly clinical potential miRNAs (miR-23c, miR-421, miR-148a, miR-676) in HCC for the further study (Fig. 3B). Anti-Argonaute 2 (AGO2) complex RNA immunoprecipitation (RIP) assays were routinely used to purify the interactive miRNAs ^20^. By using Anti-AGO2 complex RIP assays, we confirmed that AGO2 could accumulate circMRPS35 and these 4 miRNAs candidates (Fig. 3C and D, S2C). However, when overexpressed this circMRPS35 (Fig. S2B), we observed that only miR-148a was significantly accumulated than rest of other 3 miRNAs, which suggested that miR-148a was associated with circMRPS35 both in Huh-7 and HCC-LM3 cells (Fig. 3C and D, S2C). Furthermore, we found that the expression of miR-148a was significantly decreased in both HCC tissues (n=35) and 5 HCC cell lines (Fig. 3E and F).

**Figure 3.**
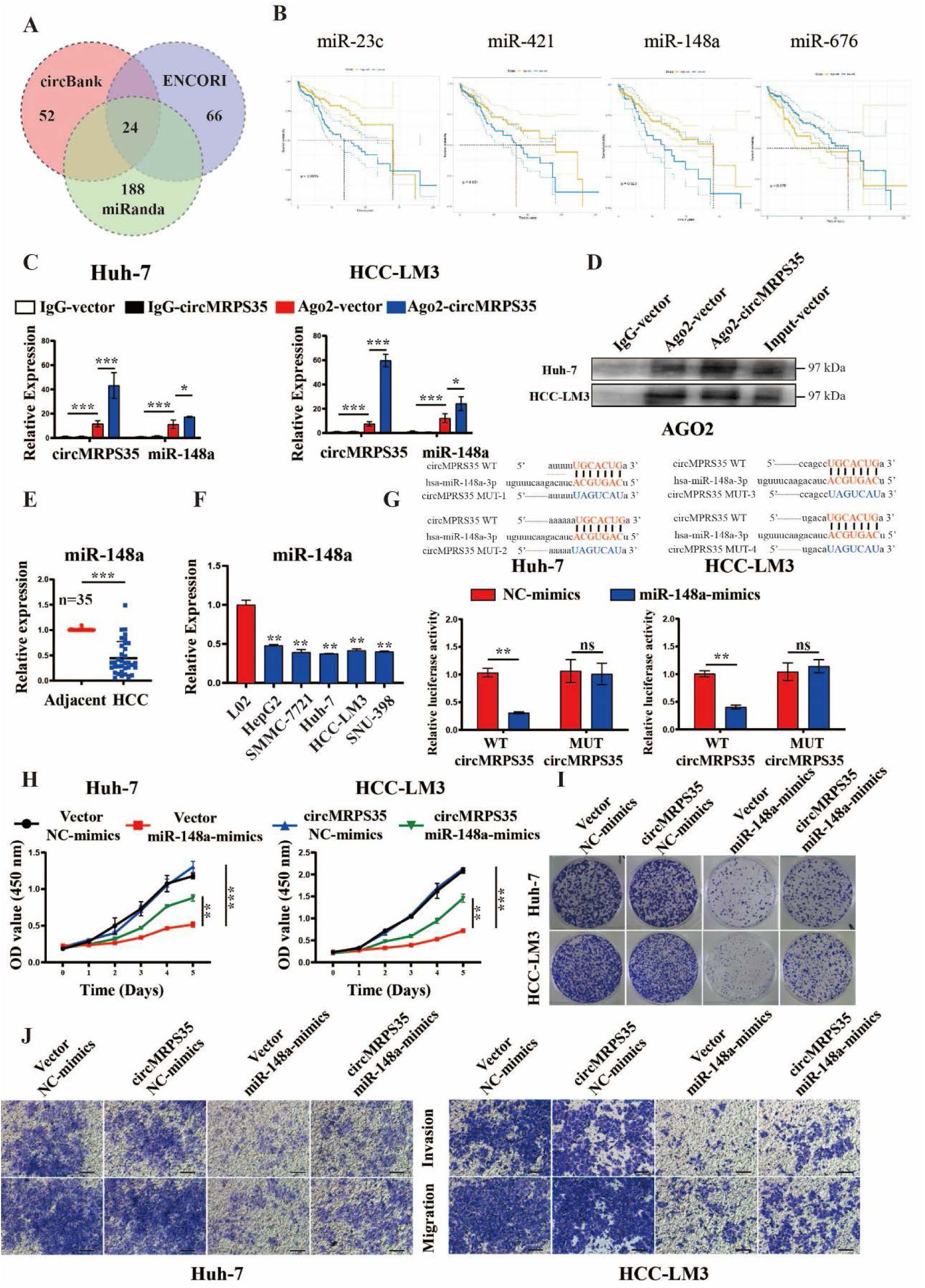
CircMRPS35 serves as a sponge for miR-148a in HCC cells. (A) Schematic illustration of the target miRNAs of circMRPS35 predicted by overlapping miRanda, ENCORI and circBANK database. (B) Kaplan-Meier analysis of the miR-23c, miR-421, miR-148a, miR-676 in HCC. (C) RT-qPCR analysis of circMRPS35 and miR-148a with AGO2-RIP. (D) Western blot analysis of AGO2 protein level in Huh-7 and HCC-LM3 cells. (E) RT-qPCR analysis of miR-148a in 35 pairs of HCC and adjacent tissues. (F) RT-qPCR analysis of miR-148a in HCC cell lines compared to L02. (G) Predicted complementary binding sites between circMRPS35 and miR-148a (up), and luciferase reporter assay was used to test the binding of miR-148a and circMRPS35 in Huh-7 and HCC-LM3 cells (down). (H-J) Co-transfection with miR-148a mimics and circMRPS35 to test the proliferation assays (H), colony formation assays (I), migration and invasion assays (J) in Huh-7 and HCC-LM3 cells. Error bars represent the means ± SEM of 3 independent experiments. \**P* < 0.05, \*\**P* < 0.01, \*\*\**P* < 0.001.

Notably, by analyzing Target Scan database, we found that circMRPS35 had 4 binding sites with miR-148a (Fig. S2D). Dual-luciferase reporter system was used to detect the interaction between circMRPS35 and miR-148a. We observed that miR-148a inhibited the relative luciferase intensity of circMRPS35 contained luciferase vector, compared with the 4 sites mutant vector in Huh-7 and HCC-LM3 cells, respectively (Fig. 3G, S2E).

In addition, a series rescue assays were carried out to investigate the regulation of circMRPS35-miR-148a axis in HCC progression. Results from the cell proliferation, clone formation, migration and invasion assays corroborated that the restraining influence of miR-148a was reversed by the stable circMRPS35 overexpression in Huh-7 and HCC-LM3 cells (Fig. 3H-J).

Overall, these results provided the solid evidence that the oncogenic functions of circMRPS35 were acted through sponging miR-148a in HCC.

### CircMRPS35 sponges miR-148a and in turn regulates STX3-PTEN axis in HCC cells

By using multiple databases (TargetScan, MirWork, MirDB and TCGA), we investigated the downstream targets of circMRPS35-miR-148a axis and screened out 5 genes, including Syntaxin 3 (*STX3*), Leptin receptor overlapping transcript like 1 (*LEPROTL1*), Macrophage immunometabolism regulator (*MACIR*), Tyrosine 3-monooxygenase/tryptophan 5-monooxygenase activation protein beta (*YWHAB*), and Ubiquitin conjugating enzyme E2 D1 (*UBE2D1*), which were significant negatively correlated with the expression of miR-148a in HCC (Fig. 4A and B, S3A-D). Further studies showed that these 5 genes were highly expressed in HCC cells (Fig. 4C, S3E-H). However, only *STX3* was markedly regulated by miR-148a in Huh-7 and HCC-LM3 cells, and higher STX3 expression had worse prognosis in patients (Fig. 4D and E, S3I-L). Further, we confirmed that STX3 was highly expressed in HCC tissues (Fig. 4F and G). In addition, we found that miR-148a mimic decreased the relative luciferase intensity of STX3’ 3’-untranslated region (3’-UTR) contained luciferase vector, compared to the mutant vector in Huh-7 and HCC-LM3 cells by the dual-luciferase reporter assay (Fig. 4H), respectively.

**Figure 4.**
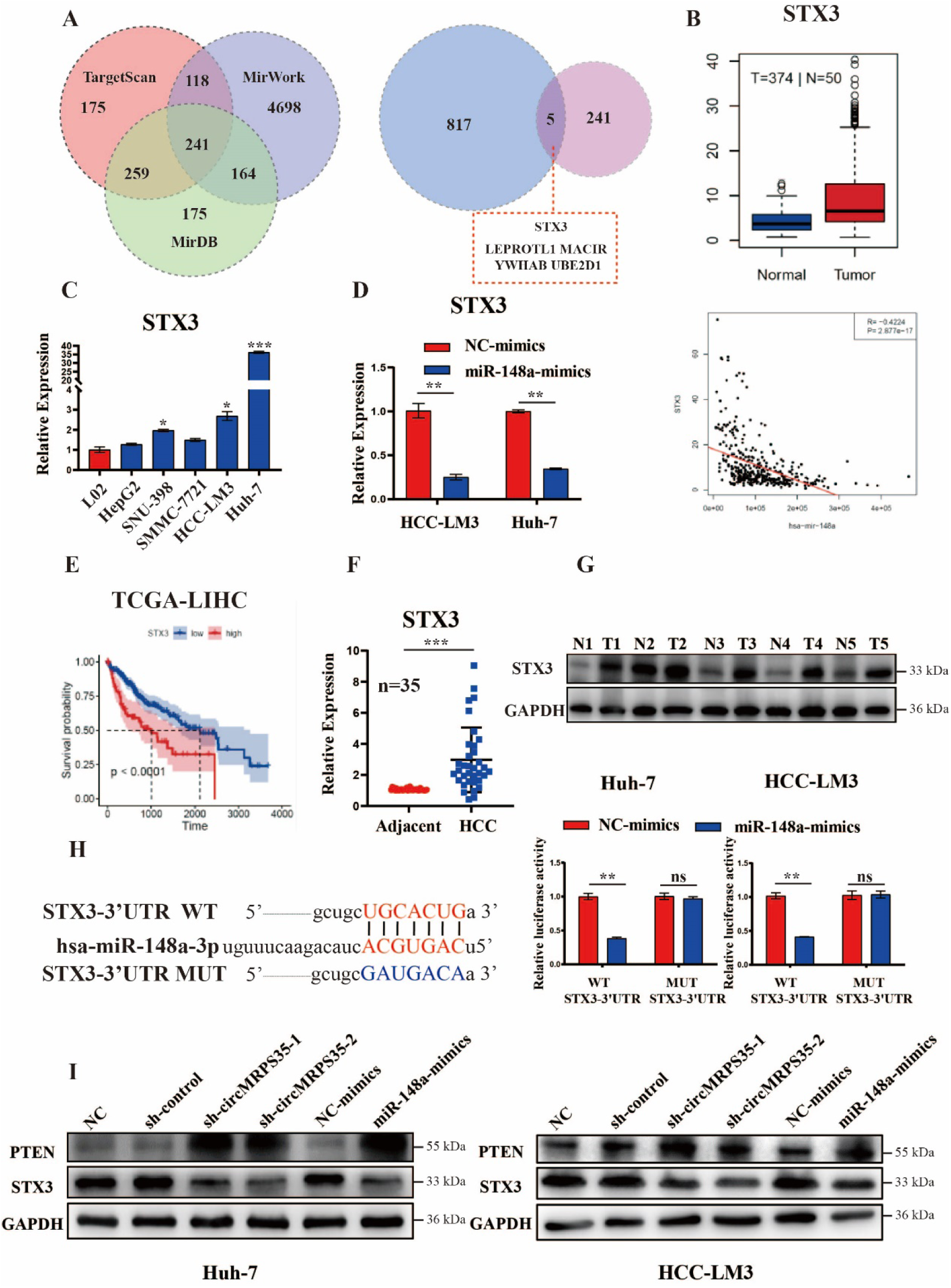
CircMRPS35 sponges miR-148a and in turn regulates STX3-PTEN axis in HCC cells. (A) Schematic illustration showing the target mRNAs of miR-148a predicted by overlapping TargetScan, MirWork and MirDB database (left) and HCC TCGA database (right). (B) TCGA analysis of expression of STX3 in HCC tissues and correlation analysis of miR-148a and STX3 expression. (C) RT-qPCR assays of STX3 expression in HCC cell lines compared to L02 cells. (D) RT-qPCR assays of STX3 in miR-148a overexpression HCC-LM3 and Huh-7 cells. (E) Kaplan-Meier analysis of the expression of STX3 in HCC. (F) RT-qPCR analysis of STX3 in 35 pairs of HCC and adjacent tissues. (G) Western blot analysis of STX3 in 5 pairs of HCC and adjacent tissues. (H) Predicted complementary binding sites between STX3 and miR-148a (left), and luciferase reporter assay was used to test the binding of STX3 and miR-148a in Huh-7 and HCC-LM3 cells (right). (I) Western blot analysis of STX3 and PTEN in silenced circMRPS35 and miR-148a overexpression and control HCC-LM3 and Huh-7 cells. Error bars represent the means ± SEM of 3 independent experiments. \**P* < 0.05, \*\**P* < 0.01, \*\*\**P* < 0.001.

Previous study found that STX3 could degrade the phosphatase and tensin homolog (PTEN) by increasing its ubiquitination, thus resulting in activation of the PI3K-Akt-mTOR signaling ^21^. We further observed that STX3 was downregulated in stable circMRPS35 silenced and miR-148a overexpressed Huh-7 and HCC-LM3 cells (Fig. 4I). In contrast, PTEN was upregulated both in stable circMRPS35 silenced and miR-148a overexpressed Huh-7 and HCC-LM3 cells (Fig. 4I).

Overall, we demonstrated that circMRPS35 regulated STX3-PTEN axis in HCC cells through sponging miR-148a.

### Chemotherapy induces the expression of circMRPS35 and translation of circMRPS35-168aa

By further re-analyzing the RNA-seq database (GSE140202), we found that circMRPS35 was highly expressed in Sorafenib treated group, compared to the none-treated group in HCC (Fig. 5A), which indicated that circMRPS35 might be related to chemotherapy.

**Figure 5.**
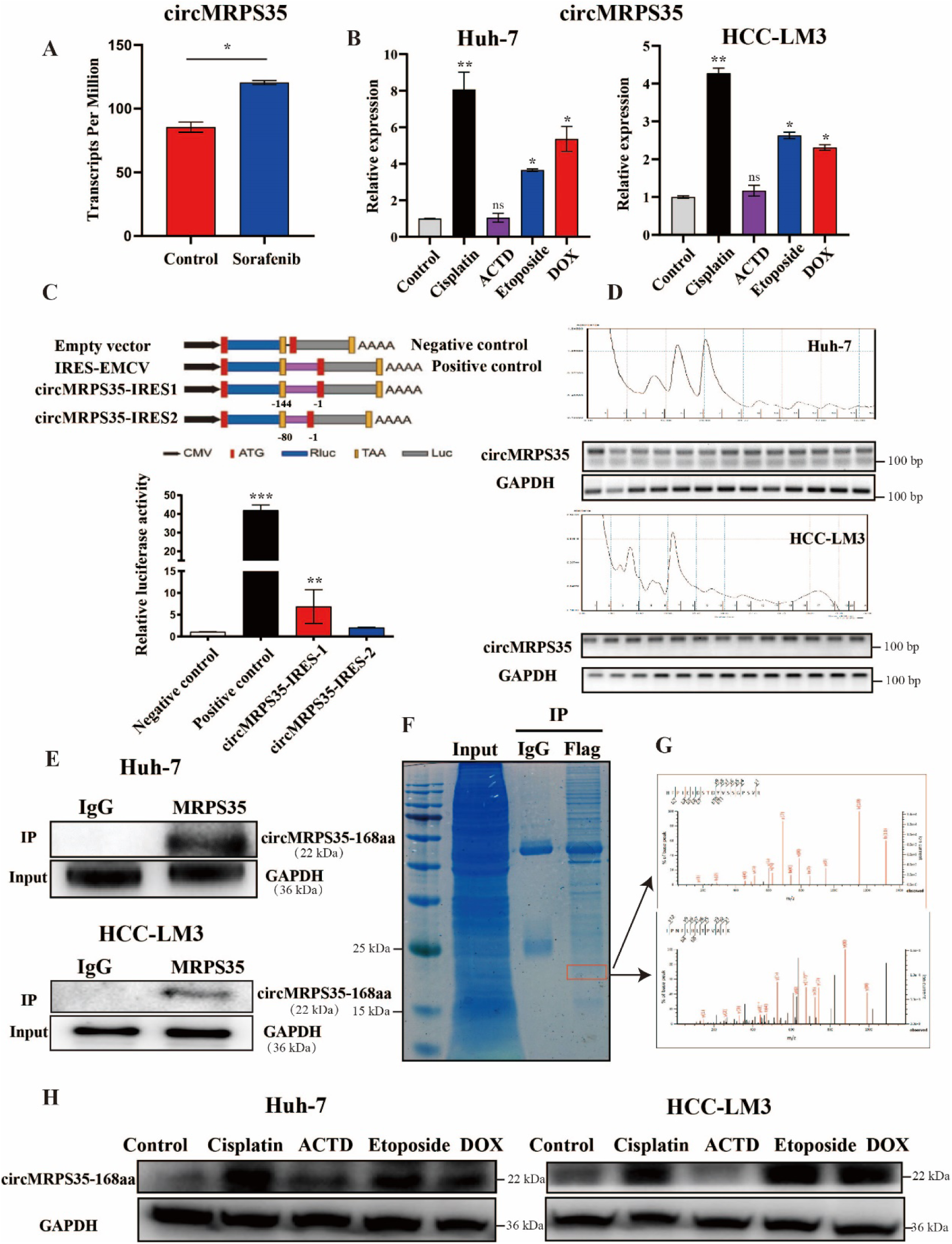
Chemotherapy induces the expression of circMRPS35 and translation of circMRPS35-168aa. (A) Transcripts pre million analysis of circMRPS35 in Sorafenib treatment cells compared to none-treatment cells by RNA-seq (GSE140202). (B) RT-qPCR analysis of circMRPS35 after DOX, Etoposide, ACTD and cisplatin treatment or none-treated Huh-7 and HCC-LM3 cells. (C) Schematic illustration showing that IRES sequences in circMRPS35 were cloned between Rluc and Luc reporter genes with independent start and stop codons (up). The relative luciferase activity of Luc/ Rluc in the above vectors was tested in Huh-7 cells (down). Encephalomyocarditis Virus (EMCV) IRES was used as a positive control. (D) Polysome fractionation and RT-PCR analysis of Huh-7 and HCC-LM3 cell lysate, and GAPDH as the positive control. (E) IP by MRPS35 antibody and western blot assay of circMRPS35-168aa in Huh-7 and HCC-LM3 cells, GAPDH as the positive control. (F-G) IP by Flag antibody and SDS-PAGE separation of protein bands stained by Coomassie brilliant blue (CBB) and the band (red frame) (left) analyzed by LC-MS (right). (H) Western blot analysis of the expression of circMRPS35-168aa after 4 chemotherapy drugs treatment in Huh-7 and HCC-LM3 cells and GAPDH as the positive control. Error bars represent the means ± SEM of 3 independent experiments. \**P* < 0.05, \*\**P* < 0.01, \*\*\**P* < 0.001.

To verify whether circMRPS35 was induced by multiple chemotherapeutic drugs’ treatment, we then used other 4 commonly used chemotherapeutic drugs (Doxorubicin (DOX), ACTD, Etoposide and cisplatin) to treat Huh-7 and HCC-LM3 cells and found that the expression of circMRPS35 was highly elevated in 3 chemotherapeutic drugs (DOX, Etoposide and cisplatin), and the most highly expression of circMRPS35 was induced by cisplatin, compared to the none-treated cells (Fig. 5B). Therefore, we used cisplatin for further studies. In this study, we had showed that stable circMRPS35 overexpression did not promote malignant progression in Huh-7 and HCC-LM3 cells compared to the control cells (Fig. 3H-J), and no significantly different expressions of *STX3* (the downstream of circMRPS35) and miR-148a were observed among groups (Fig. S4A and B), which revealed that the elevated expression of circMRPS35 might have other functions in HCC cells rather than through sponging miR148a in cisplatin treatment.

A recent study has showed that circMRPS35 serves as a protein binding RNA for the transcriptional activation of Forkhead box O1 (*FOXO1*) and Forkhead box O3a (*FOXO3a*) in gastric cancer ^22^. However, we did not find the different expressions of *FOXO3a* and *FOXO1* in cisplatin treated HCC-LM3 and Huh-7 cells compared to the none-treated cells (Fig. S4C and D), therefore circMRPS35 did not serve as a protein binding RNA to regulate the expression of *FOXO1* and *FOXO3a* in cisplatin treated HCC cells. Based on the above results, we hypothesized that there were other functions of circMRPS35 in the condition of cisplatin treatment.

Few studies had shown that translation of some circRNAs could occur through IRES ^23, 24^. By analysis from circRNADb database, we found that circMRPS35 had two putative internal ribosome entry site (IRES) regions (14-158 sites and 81-161 sites) with a crossing back-spliced sites open reading frame (ORF), which potentially codes a 168 amino-acid peptide (Fig. S4E).

To examine the putative IRES activity in circMRPS35, we used a modified dual luciferase reporter system (the promoter of firefly luciferase was removed) and obtained that the IRES (14-158 sites) induced the high F-Luc/R-Luc activity compared to the truncated IRES (81-161 sites) (Fig. 5C). This result suggested that the activity of IRES (14-158 sites) of circMRPS35 do induce the translation of its ORF.

The translation process of circRNAs could be associated with polyribosome (polysome) ^25, 26^. Furthermore, separation of polysome fractionation was used to detect the circMRPS35 distribution. The results showed that circMRPS35 was present in all fractions including monosome and polysome fractions in HCC-LM3 and Huh-7 cells (Fig. 5D).

Then, we detected the endogenous translational capacity of circMRPS35. By analysis the sequence of this peptide, we found that 115 amino-acids of this peptide were originated from MRPS35 and the rest of 53 amino-acids was unique. By using the immunoprecipitation of MRPS35 antibody to detect this peptide of circMRPS35 (circMRPS35-168aa), a 22-kDa band was identified by Western blot in Huh-7 and HCC-LM3 (Fig. 5E). Then we used immunoprecipitation (IP) of flag antibody in circMRPS35-flag overexpressed Huh-7 cells, and this 22 kDa band was further detected and identified by Liquid chromatograph-mass spectrometer (LC-MS) (Fig. 5F and G). We thus confirmed that this protein was circMRPS35-168aa with the identified short amino acid sequences (Fig. 5G). By treating Huh-7 and HCC-LM3 cells with the 4 commonly used chemotherapeutic drugs we found that this circMRPS35-168aa was significantly induced by DOX, Etoposide and cisplatin (Fig. 5H).

Taken together, we demonstrated that circMRPS35 contemporarily encoded an uncharacterized peptide induced by multiple chemotherapeutic drugs in HCC cells.

### circMRPS35-168aa resists the cisplatin treatment in HCC cells

To further confirm the relationship between circMRPS35-168aa and chemotherapy, we used these chemotherapeutic drugs to identify the most sensitive drug regulated by circMRPS35-168aa. Western blot analysis firstly ensured that circMRPS35-168aa was stably overexpressed in Huh-7 and HCC-LM3 cells (Fig. S4F).

Cell viability analysis showed that the overexpression of circMRPS35-168aa mostly induced cisplatin resistance, while low expression of circMRPS35 mostly inhibited the cell growth with cisplatin treatment compared to DOX, ACTD and Etoposide both in Huh-7 and HCC-LM3 cells (Fig. 6A-C, S4G).

**Figure 6.**
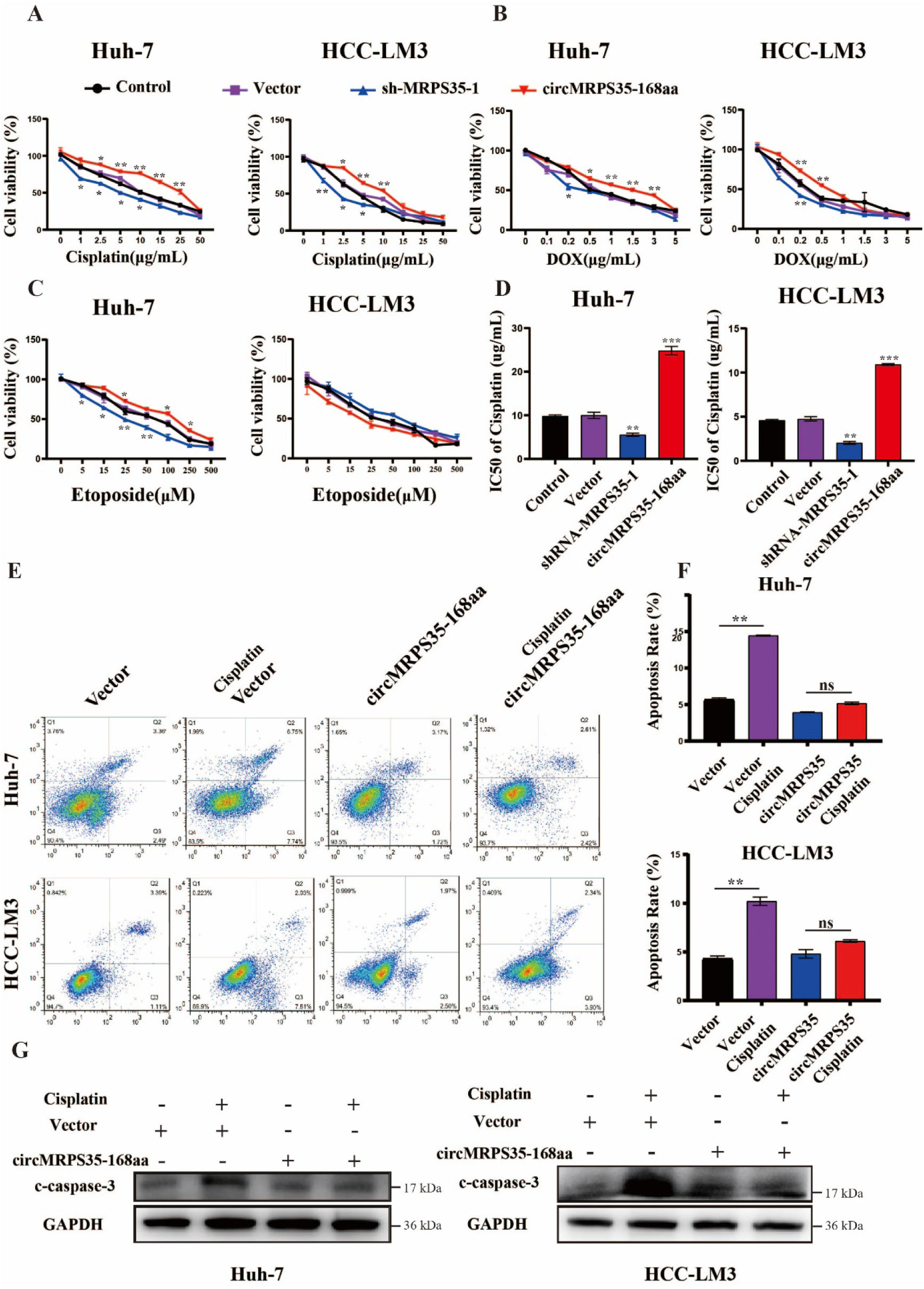
circMRPS35-168aa resists the cisplatin treatment in HCC cells. (A) Cell viability assay of different circMRPS35-168aa expressed Huh-7 and HCC-LM3 cells with different concentrations of cisplatin treatment. (B) Cell viability assay of different circMRPS35-168aa expressed Huh-7 and HCC-LM3 cells with different concentrations of DOX treatment. (C) Cell viability assay of different circMRPS35-168aa expressed Huh-7 and HCC-LM3 cells with different concentrations of Etoposide treatment. (D) IC50 analysis of different circMRPS35-168aa expressed Huh-7 and HCC-LM3 cells with different concentrations of cisplatin treatment. (E-F) FACS analysis of apoptosis about different circMRPS35-168aa expressed Huh-7 and HCC-LM3 cells with cisplatin treatment (left), and statistical analysis of apoptosis rate (right). (G) Western blot analysis of cleaved Caspase3 in different circMRPS35-168aa expressed Huh-7 and HCC-LM3 cells with cisplatin treatment, and GAPDH as the positive control. Error bars represent the means ± SEM of 3 independent experiments. \**P* < 0.05, \*\**P* < 0.01, \*\*\**P* < 0.001.

The half maximal inhibitory concentration (IC50) was also decreased in low circMRPS35 expressed Huh-7 and HCC-LM3 cells and increased in circMRPS35 overexpressed Huh-7 and HCC-LM3 cells with cisplatin treatment (Fig. 6D). Apoptosis analysis showed that the apoptosis rate was decreased in circMRPS35-168aa overexpressed Huh-7 and HCC-LM3 cells with cisplatin treatment (Fig. 6E and F). Western blot results showed that the high expression of circMRPS35-168aa counteracted cisplatin induced high level of the cleaved Caspase-3 (c-Caspase-3) (Fig. 6G).

In summary, our results showed that circMRPS35-168aa can play a critical role in cisplatin resistance in HCC cells.

## Discussion

Circular RNAs (circRNAs), characterized by high stability and conservation, have been increasingly demonstrated to function as the novel promising therapeutic RNA molecules for diverse human diseases, including cancers ^27^. Previous studies had shown that several circRNAs correlated with pathogenesis, clinical pathological and prognostic for HCC diagnosis ^28, 29^. However, a conclusive or practical criterion for the HCC diagnosis still requires further study ^30, 31^. In this study, by using the clinical RNA-seq databases, HCC cell lines, human HCC tissues and the HCC xenograft mouse models, we found that circMRPS35 was significantly upregulated in HCC, and stable silenced expression of circMRPS35 suppressed the growth and migration of HCC cells. Surprisingly, we demonstrated that a novel peptide encoded by circMRPS35 (circMRPS35-168aa), which was significantly induced by chemotherapeutic drugs, and promoted cisplatin resistance in HCC.

circRNAs have multiple functions ^32, 33^. Conventionally, most studies showed that circRNAs acted as the sponges of miRNAs to regulate the downstream gene expressions. Han *et al.* showed that circMTO1 acted as an endogenous sponge for miR-9 to regulate the progression of HCC ^12^. Hu *et al.* showed that circASAP1 sponged miR-326/miR-532-5p to control the MAPK1/CSF-1 signaling in HCC ^34^. Recently, other novel functions of circRNAs were reported in HCC. Zhu *et al.* found that circZKSCAN1 suppressed the transcriptional activity of Wnt/β-catenin signal pathway through competitively binding to fragile X mental retardation protein (FMRP) in HCC^13^. Liang *et al.* identified a coding circRNA derived from β-catenin, which could activate Wnt/β-catenin pathway to promote the progression of HCC ^14^. However, in this study, we found that circMRPS35, on the one hand, could act as miRNA sponge, forming a circMRPS35-miR148a-STX3-PTEN axis to control malignant progression of HCC cells. On the other hand, circMRPS35 could encode a novel 168 amino-acid peptide endowing the HCC cells with chemoresistance in chemotherapeutic drugs treatment.

A previous study has shown that circMRPS35 was low expressed and acted as a protein sponge of Lysine acetyltransferase 7 (KAT7) for histone acetylation to regulate the transcriptions of FOXO1 and FOXO3a in gastric cancer ^22^. In contrast to this gastric cancer study, we found that circMRPS35 was highly expressed and with other multiple functions in HCC rather than as a KAT7 sponge, and we did not find the different expressions of *FOXO1* and *FOXO3a* in cisplatin treated HCC cells. This discrepancy may be due to the complicated roles of circMRPS35 in various cancers, and the actions of circMRPS35 may depend on the context of its binding targets inside the particular cells, or under various conditions. Why does circMRPS35 in different tissues have different functions and what factors regulate its expression and functions need to be further studied.

The increased cisplatin chemoresistance is the main problem of HCC chemotherapy, however, the mechanism of cisplatin chemoresistance remains unclear ^35–37^. A study has shown that circRNA_101505 was downregulated in cisplatin-resistant HCC tissues and circRNA_101505 could increase the sensitivity to cisplatin in HCC cells by sponging miR-103 ^38^. Another study showed that circ_0003418 suppressed tumorigenesis and cisplatin in HCC through regulating Wnt/β-Catenin pathway ^39^. Differing from those studies, we found that circMRPS35 was highly induced by cisplatin, and which coded a circMRPS35-168aa to resist cisplatin treatment in HCC cells. Mechanically, circMRPS35-168aa could suppress the cisplatin induced apoptosis through inhibiting the cleavage of Caspase 3 in HCC.

A few studies had showed that the expressions of circRNAs were regulated in cisplatin treatment and led to cisplatin resistant in HCC. One study has showed that the expression of circRNA_102272 was up-regulated in cisplatin treated HCC cells and promoted cisplatin resistance by sponging miR-326 to regulate RUNX2 axis ^40^. Another study has shown that the expression of circFN1 was enhanced in cisplatin-resistant gastric cancer tissues and cells and promoted cisplatin resistance via sponging miR-182-5p ^41^. Similar with those studies, we found that circMRPS35 was highly expressed in HCC, and chemotherapy further elevated its expression. Differing from these studies, we found that a circMRPS35-168aa coded by circMRPS35 directly promoted cisplatin resistance in HCC. However, the regulating mechanisms of circMRPS35 expression pattern under different conditions is still unknown. In further study, we are going to find the interacting proteins of circMRPS35-168aa for mechanical studies of cisplatin resistance in HCC. In addition, our study might put forward a new insight for selections of therapeutic drugs, which not only inhibited the malignant progression, but also suppressed chemotherapy resistance in cancers’ treatment.

In current study, the clinical evidence of circMRPS35 were still limited and the correlation between circMRPS35 and clinical diagnosis, prognostic, pathogenesis and chemoresistance of HCC needed to be further studied. In the further study, we will continue collecting HCC tissues (with or without chemotherapy) and recording the corresponding follow-up information to investigate the relation between circMRPS35 and prognostication of HCC, and we will use nude mice models to further confirm the cisplatin sensitivity in HCC cells with different levels of circMRPS35-168aa expressions.

In summary, by using functional verification together with clinical evidence, the present study demonstrated that circMRPS35 could be a crucial regulator for the progression and chemoresistance in HCC with its different expression pattern under different conditions. circMRPS35 not only elicited its oncogenic role in HCC through sponging miR-148a to regulate STX3-PTEN axis, but also further upregulated in chemotherapeutic drugs treatment which stimulated the coding of circMRPS35-168aa peptide. circMRPS35-168aa suppressed the cisplatin induced apoptosis through inhibiting the cleavage of Caspase3, which led to cisplatin resistance (Fig. 7). Taken together, we provided that circMRPS35 has the potential to be a biomarker to predict prognosis for HCC therapy and a therapeutic target for HCC, especially in HCC chemoresistance.

**Figure 7.**
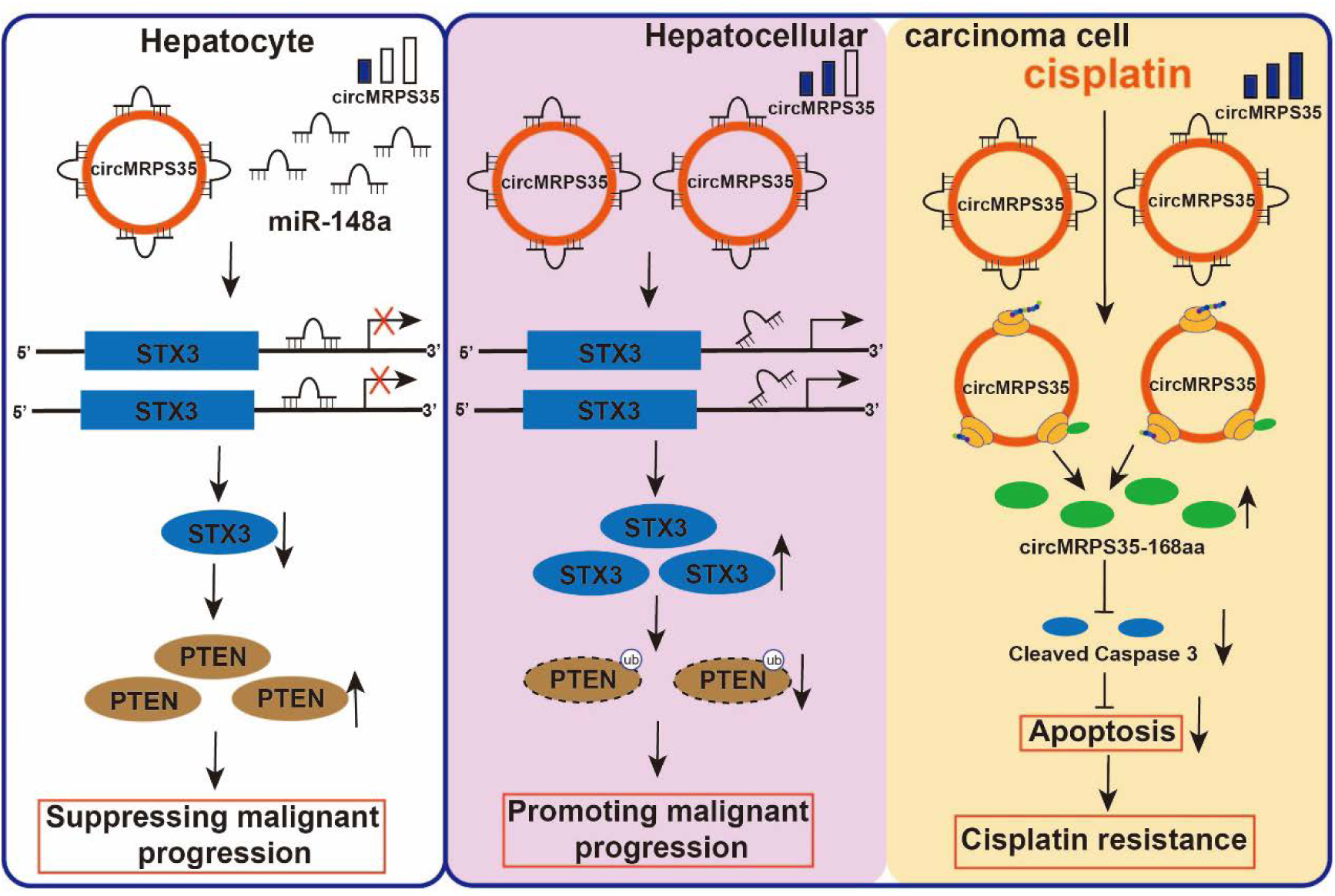
Diagram models of the effects about circMRPS35 HCC. In this model, circMRPS35 elicited its oncogenic role in HCC through sponging miR-148a to regulate STX3-PTEN axis, and circMRPS35 further upregulated in chemotherapeutic drugs treatment which stimulated the coding of circMRPS35-168aa peptide. circMRPS35-168aa suppressed the cisplatin induced apoptosis through inhibiting the cleavage of Caspase3, which led to cisplatin resistance.

## Materials and methods

### Patients and tissue samples

In this study, 35 pairs of HCC and their corresponding adjacent tissues were collected and stored at −80°C from patients who underwent surgery at Chinese PLA General Hospital between 2018 and 2020. None of the patients was treated with either chemotherapy or radiation prior to surgery. Clinical data of patients were summarized in Table 1.

**Table 1.** Association between circMRPS35 expression and clinical features of HCC.

| Clinical features | *n* | High expression | Low expression | *P* value |
| --- | --- | --- | --- | --- |
| Gender |  |  |  |  |
| Male | 29 | 25 | 4 | 0.8973 |
| Female | 6 | 5 | 1 |  |
| Ages (years) |  |  |  |  |
| <50 | 16 | 14 | 2 | 0.8831 |
| >50 | 19 | 16 | 3 |  |
| Tumor size (cm) |  |  |  |  |
| <5 | 17 | 14 | 3 | 0.2858 |
| >5 | 18 | 16 | 2 |  |
| Lymph node metastasis |  |  |  |  |
| No | 25 | 21 | 4 | 0.013 |
| Yes | 10 | 9 | 1 |  |
| HBV infection |  |  |  |  |
| Yes | 31 | 27 | 4 | 0.0429 |
| No | 4 | 3 | 1 |  |
| TNM stage |  |  |  |  |
| I - II | 14 | 13 | 1 | 0.0564 |
| III - IV | 21 | 17 | 4 |  |

### Bioinformatics procedure for circRNA expression analysis

HCC RNA-seq data (GSE77509, GSE114564 and GSE159220) was downloaded from the NCBI SRA database. CIRI2, CIRCexplorer2, and find_circ were used for characterization of circRNAs ^42^. HISAT2, Bowtie2 and StringTie were performed to re-assemble the sequencing transcriptome after aligning to reference genome Human GRCh37. Then, the quantification of these circRNAs was performed by using a modified version of edgeR in CIRIquant, and circBase was used for annotation of these circRNAs. The differentially expressed circRNAs were identified by using the edgeR package (version 3.12.1) with general linear model, and fold change > 2 and *P* value < 0.05 were recognized as significantly differentially expressed circRNAs.

### Cell culture

The human 293T cell, human HCC cell lines HepG2, SNU-398, SMMC-7721, Huh-7, HCC-LM3 and the human normal liver cell line L02 were used in the present study. The cell lines of 293T and HepG2 were purchased from the Cell Bank of the Peking Union Medical College Hospital (China). Rest of other cell lines were a generous gift from State Key Laboratory of Proteomics, Beijing Proteome Research Center, Beijing Institute of Radiation Medicine (China). L02, SNU-398 cell lines were cultured in Roswell Park Memorial Institute 1640 medium (Invitrogen, USA), and other cell lines were cultured in Dulbecco’s modified Eagle’s medium (Invitrogen, USA) with 10% Foetal Bovine Serum (GIBCO, Brazil) at 37°C with 5% CO_2_.

### RNA extraction and reverse transcription

Total RNAs were extracted from cell lines and tissues using Trizol (Invitrogen, USA) according to the manufacturer’s instructions. cDNAs were synthesized from total RNA using Moloney Murine Leukemia Virus (M-MLV) Reverse Transcriptase (Takara, Japan) based on the manufacturer’s instructions.

### RNase R treatment and actinomycin D assay

Total RNAs was treated with RNase R for 30 min at 37°C using 3 U/mg of RNase R (Lucigen, USA). HCC-LM3 and Huh-7 cells treated with actinomycin D (1 µg/mL) (ACTD, Sigma, USA) at 0 h, 2 h, 6 h, 12 h and 24 h before RNA extraction.

### Nucleocytoplasmic separation

The RNA of nuclear and cytoplasmic was separated and extracted using PARIS Kit (Life technologies, USA) according to the manufacturer’s instructions.

### RT-PCR and RT-qPCR

RT-PCR was conducted using PrimeSTAR Master Mix (Takara, Japan) according to manufacturer’s instructions along with PCR control. Products were separated on a 2% agarose gel and visualized with GelRed (Beyotime, China). RT-qPCR analyses were performed by using SYBR Green PCR Master Mix (Applied Biosystems, USA) with the StepOnePlus System (Applied Biosystems, USA) according to manufacturer’s instructions. GAPDH or U6 was used as the internal control, and the relative expression of target genes was calculated by 2^-ΔΔCt^ method. Primers are listed in Table S1.

### Oligonucleotide synthesis, plasmid construction and transfection

The oligonucleotides of miR-148a mimics, and control mimics were synthesized by GenePharma (China). Two specific shRNAs for circMRPS35 designed to target the covalent closed junction were cloned into PLKO.1-TRC plasmid to silence the expression of circMRPS35. The PLO5-ciR plasmid (GENESEED, China) containing the sequence of circMRPS35 was constructed and used to upregulate circMRPS35 expression. The PLV plasmid containing the sequence of circMRPS35-168aa was constructed and used to upregulate circMRPS35-168aa expression. For Dual-luciferase reporter gene assay, wild type (WT) and mutant (Mut) of miR-148a putative binding sites reporter plasmids were constructed using the circMRPS35 and 3’-UTR of STX3 sequences in the psiCHECK2 vector (Promega, USA). For IRES activity analysis, the promoter region of Renilla luciferase in psiCHECK2 vector was deleted and the IRES sequence was cloned behind the firefly luciferase. Plasmids, miR-148a mimics, and the negative controls were transfected into cells by using Lipofectamine 3000 (Invitrogen, USA) based on the manufacturer’s instructions.

### Lentivirus packaging, infection and puromycin selection

Lentiviral vectors were co-transfected with packaging plasmids psPAX2 and pMD2.G (Addgene, USA) into 293T cells. Infectious supernatant was harvested at 48 and 72 h after transfection, and filtered through 0.45 µm filters (Millipore, USA). Cells were infected by recombinant lentivirus for 48 h and then selected by appropriate concentration of puromycin for 72 h.

### Cell proliferation assays and wound healing assay

Huh-7 and HCC-LM3 cells reseeded in 96-well plates (1 × 10^3^ cells per well), and the cell viability was detected by cell counting kit-8 (CCK-8, Beyotime, China) with absorbance of wavelength of 450 nm for each well. For cell colony formation assays, the treated Huh-7 and HCC-LM3 were placed in 6-well plates (3 × 10^3^ cells per well) incubated at 37°C with 5% of CO_2_ for 7 days. Cells were stained with Crystal Violet Staining Solution (Beyotime, China). For wound healing assay the treated Huh-7 and HCC-LM3 from different groups were placed in 6-well plates (4 × 10^4^ cells per well) with serum-free medium. Constant diameter strips were scratched in the confluent monolayers with a 10 µL sterile Eppendorf pipette tip. The width of scratches was obtained at 0 and 48 h in same places using the microscope (Ti-U, Nikon, Japan).

### Migration and invasion assay

Transwell was used for invasion and migration assays. For migration assays, Huh-7 and HCC-LM3 cells reseeded in the small chambers (2 × 10^4^ cells per well), and 600 µL of cell culture medium added in the bottom chambers at 37°C with 5% of CO2 for 48h. For invasion assays, firstly, the small chambers were coated with 100µL Matrigel for 30 min incubation in 37°C, and then Huh-7 and HCC-LM3 cells reseeded in the small chambers (2 × 10^4^ cells per well), and 600 µL of cell culture medium added in the bottom chambers at 37°C with 5% of CO_2_ for 48h. Cells were stained with Crystal Violet Staining Solution (Beyotime, China), and removed inner cells of small chambers. The cells of outer cells were photographed randomly by the microscopy (Ti-U, Nikon, Japan).

### Cell cycle analysis

Treated Huh-7 and HCC-LM3 cells (2 × 10^5^ cells) were digested by trypsin, washed twice with PBS, and fixed 4 h at 4°C in 70% ethanol. Cells were washed with PBS and strained with Cell Cycle Analysis Kit (Beyotime, China). Flow cytometry (BD, USA) was used to analyze the staining and the data were analyzed with FlowJo 7.6 software (USA).

### Dual-luciferase reporter gene assay

Huh-7 and HCC-LM3 cells were co-transfected with WT or Mut circMRPS35/STX3 3′-UTR and miR-148a mimics or mimics-NC using Lipofectamine 3000. Renilla luciferase activity was normalized to firefly luciferase activity. For IRES activity analysis, 293T cells was transfected with IRES contained plasmids. Firefly luciferase activity was normalized to Renilla luciferase activity. After transfection for 48 hours, cells were subjected to dual-luciferase analysis. Luciferase activity was assessed using the dual-luciferase reporter kit (TransGene, China) and performed via a dual-luciferase reporter assay system (Promega, USA).

### RIP assay

RIP assays were performed using the Magna RIP RNA-Binding Protein Immunoprecipitation Kit (Millipore, USA) with the mouse anti-Ago2 antibody (Millipore, USA) according to the manufacturer’s instructions. Mouse anti-IgG antibody (Millipore, USA) was used as a negative control.

### Western blot and IP assay

For Western blot assay, total protein of treated Huh-7 and HCC-LM3 cells was extracted by protein lysis buffer, separated by 10% SDS-PAGE gel, and transferred onto the polyvinylidene fluoride (PVDF) membrane (Millipore, USA). After the membrane was incubated with a primary antibody and corresponding secondary antibody, chemiluminescent reagent was used for detecting the signal. For IP assay, the primary antibodies were incubated with protein A/G magnetic beads (Thermo Scientific, USA) at 4°C with gentle rotation for 3 h. Lysis was incubated with the beads for 2 h at 25°C, and the precipitated complex was subjected to Western blot analysis.

### In vivo xenograft assay

Four-week-old female BALB/c nude (nu/nu) mice were purchased from the Si Pei Fu (China). Mice were housed under Specified Pathogen Free (SFP) conditions. Huh-7 and HCC-LM3 cells (2 × 10^6^ cells) with different expression of circMRPS35 were subcutaneously injected in BALB/c nude mice respectively. Tumor volumes were measured every 5 days and calculated using: volume (mm^3^) = length × width^2^/2. Tumor weights were weighed 25 days after injection.

### IHC assay

For immunostaining, sections were pretreated with hydrogen peroxide (3%) for 10 min to remove the endogenous peroxidase, followed by antigen retrieval in a microwave for 15 min in 10 mM citrate buffer (pH 6.0). Ki67 primary antibody was used at a dilution of 1:1,000 and incubated for 30 min at room temperature, followed by washing and incubation with the biotinylated secondary antibody for 30 min at room temperature and the stained with IHC Staining Kits (Boster, Beijing) according to the manufacturer’s instructions. The slides were counterstained with hematoxylin and dehydrated in alcohol and xylene before mounting. The slides were photographed randomly by the microscopy (Ti-U, Nikon, Japan).

### Polysome fractionation assay

Huh-7 and HCC-LM3 cells were pre-treated with 200 µM cycloheximide (Sigma, USA) for 5min at 37°C and washed with ice-cold PBS containing 200 µM cycloheximide. Cells were then lysed with polysome lysis buffer for 30 min on ice. After centrifugation at 14,000 rpm for 10 min at 4°C, the supernatant was loaded onto 10 mL continuous 15-50% sucrose gradients buffer containing 50 U/ml RNase inhibitor. The samples were centrifuged at 4°C for 3 h at 100,000 g by using Avanti J-30XP (Beckman, USA), and the fractions were collected using a Brandel Fractionation System (USA) and an Isco UA-6 ultraviolet detector (USA) was used to produce polysome profiles for gradients. Extraction and transcription of total RNA from each fraction and RT-PCR was conducted as showing above. GAPDH served as positive control.

### LC-MS analysis

Proteins were separated via sodium dodecyl sulfate polyacrylamide gel electrophoresis (SDS-PAGE), and gel bands were manually excised and digested with sequencing-grade trypsin (Promega, USA). The digested peptides were analyzed using a QExactive mass spectrometer (Thermo Fisher, UAS). Fragment spectra were analyzed using the National Center for Biotechnology Information nonredundant protein database with Mascot (Matrix Science, USA).

### Apoptosis analysis

Huh-7 and HCC-LM3 cells were resuspended and washed with PBS for 3 times, and cell were stained with Cell Apoptosis Analysis Kit (Beyotime, China) based on the manufacturer’s instructions. Flow cytometry (BD, USA) was used to analyze the staining and the data were analyzed with FlowJo 7.6 software (USA).

### Chemotherapeutic drugs treatment

Huh-7 and HCC-LM3 cells reseeded in 6-well plates (8 × 10^5^ cells per well) overnight, and cells were treated with 0.5 µg/mL of DOX (Sigma, USA), 50 µM of Etoposide (Sigma, USA), 5 µg/mL of cisplatin (Sigma, USA) and 0.2 µg/mL of ACTD respectively. After 24 h treatment, cells were collected for RT-qPCR and Western blot analysis.

### IC50 analysis

Huh-7 and HCC-LM3 cells reseeded in 96-well plates (5 × 10^3^ cells per well) overnight, and cells were treated with DOX (0, 0.1, 0.2, 0.5, 1, 1.5, 3, 5 µg/mL), Etoposide (0, 5, 15, 25, 50, 100, 250, 500 µM), cisplatin (0, 1, 2.5, 5, 10, 15, 25, 50 µg/mL) and ACTD (0, 0.1, 0.15, 0.2, 0.5, 1, 2.5 µg/mL) respectively for 24h. Cell viability was detected by CCK-8 kits (Beyotime, China) with absorbance of wavelength of 450 nm for each well. IC50 was ensured based on the cell viability data.

### Statistical analysis

All data are expressed as the mean ± SEM (standard error of mean). Two-tail unpaired or paired Students’ t-test was applied to analyze the differences between two groups. Data conforming to normal distribution among multiple groups were analyzed by one-way or two-way analysis of variance (ANOVA). The values of \**P* < 0.05, \*\**P* < 0.01, and \*\*\**P* < 0.001 were indicative of statistical significance and ns were indicative of nonstatistical significance. The statistical analysis was performed using GraphPad Prism 8.0. (USA).

## Acknowledgments

We thank for the instrument and equipment support provided by the platform of Institute of Nutrition and Health, China Agricultural University. We thank Prof. Gangqiao Zhou (State Key Laboratory of Proteomics, Beijing Proteome Research Center, Beijing Institute of Radiation Medicine, Beijing, China) for kindly providing the HCC cell lines.

## Author contributions

Xiangdong Li signed the project, guided experiments, and analyzed data. Peng Li, Runjie Song and Xiangdong Li interpreted the data. Peng Li and Runjie song conducted experiments. Mei Liu, Fan Yin collected the clinical data and samples. Huijiao Liu and Runjie song analyzed RNA-seq and TCGA data. Yuting Zhong, Shuoqian Ma, Xiaohui Lu and Xiaomeng Jia revised the manuscript. Xiru Li provided guidance for experiments. All authors approved the final content.

## Conflict of interest

The authors declare that they have no conflict of interest.

## Ethics Statement

The use of human tissues specimens was approved by the ethical committee of Chinese PLA General Hospital. All animal studies were approved by the ethical committee of the China Agricultural University. The study was performed in accordance with the Declaration of Helsinki.

## Funding

This study was supported by grants from the National Key Research and Development Project (2018YFC1004702), Fund of the National Natural Science Foundation of China (31970802), and Beijing Municipal Natural Science Foundation (7202099).

---

**Figure S1.**
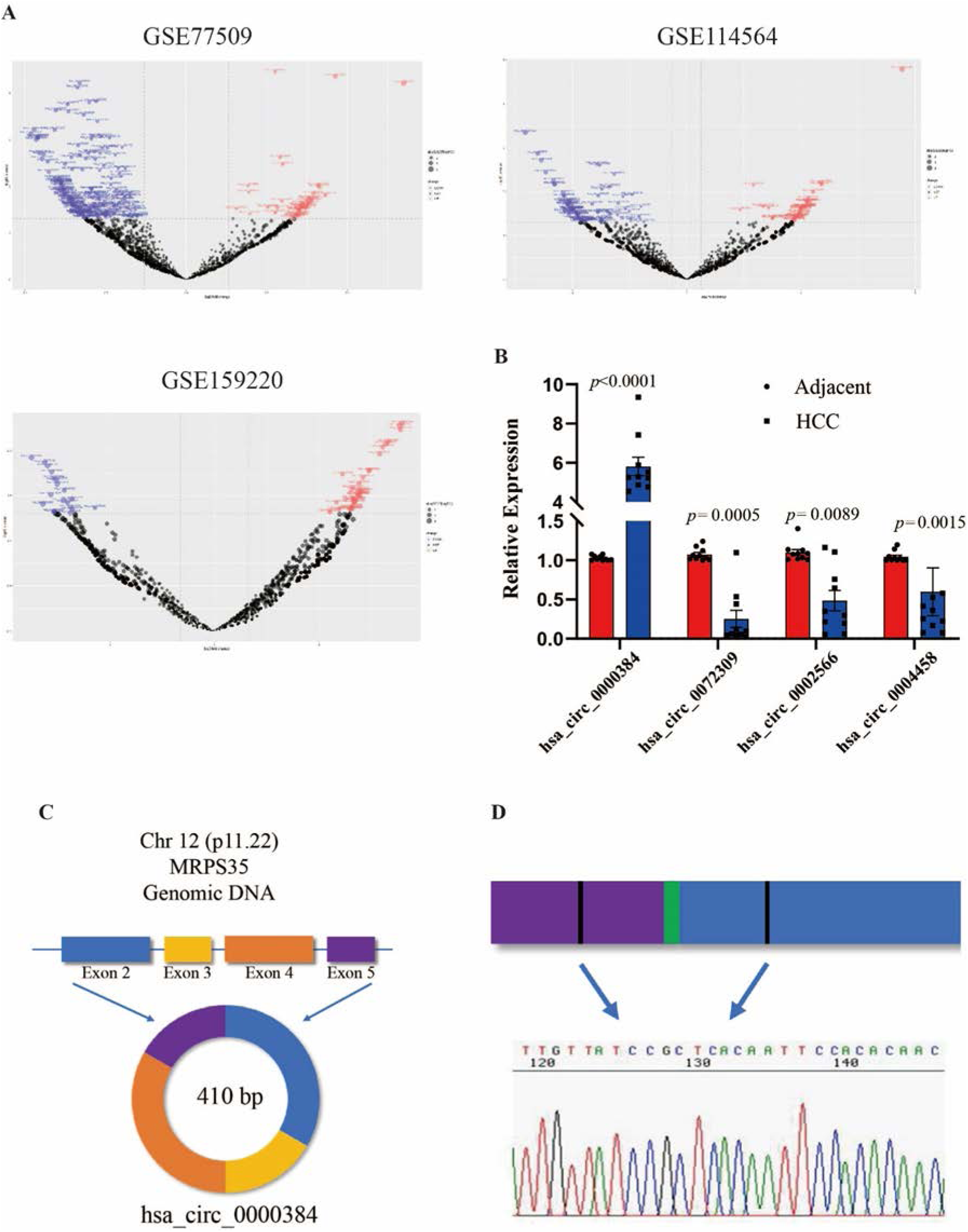
(A) Volcano plots analysis of circRNAs in 3 RNA-seq data (GSE77509, GSE114564, GSE159220). (B) RT-qPCR analysis of the 4 candidates circRNAs in HCC tissues (n=10) compared with non-tumor adjacent tissues. (C) Schematic representation of circMRPS35 formation. (D) The back-splice junction site of circMRPS35 was validated by Sanger sequencing. Error bars represent the means ± SEM of 3 independent experiments. \**P* < 0.05, \*\**P* < 0.01, \*\*\**P* < 0.001.

**Figure S2.**
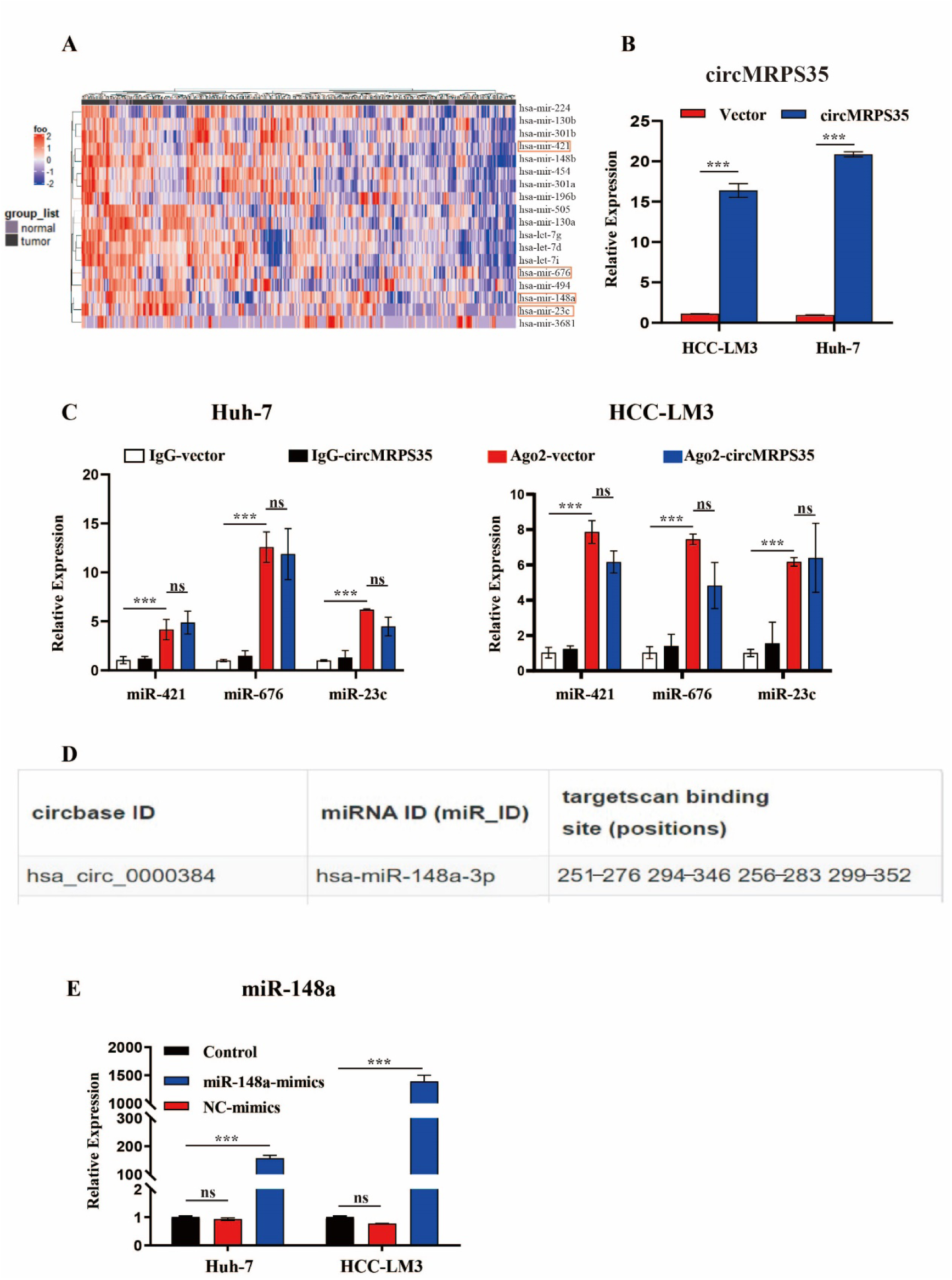
(A) Expression heat map of 24 target miRNAs with analysis of TCGA database. (B) RT-qPCR analysis of circMRPS35 in circMRPS35 overexpression Huh-7 and HCC-LM3 cells. (C) RT-qPCR analysis of circMRPS35 and miR-23c, miR-421, miR-676 with AGO2-RIP in Huh-7 and HCC-LM3 cells. (D) Binding positions of circMRPS35 and miR-148a was showed in Targetscan database. (E) RT-qPCR analysis of miR-148a in miR-148a-mimics overexpression Huh-7 and HCC-LM3 cells. Error bars represent the means ± SEM of 3 independent experiments. \**P* < 0.05, \*\**P* < 0.01, \*\*\**P* < 0.001.

**Figure S3.**
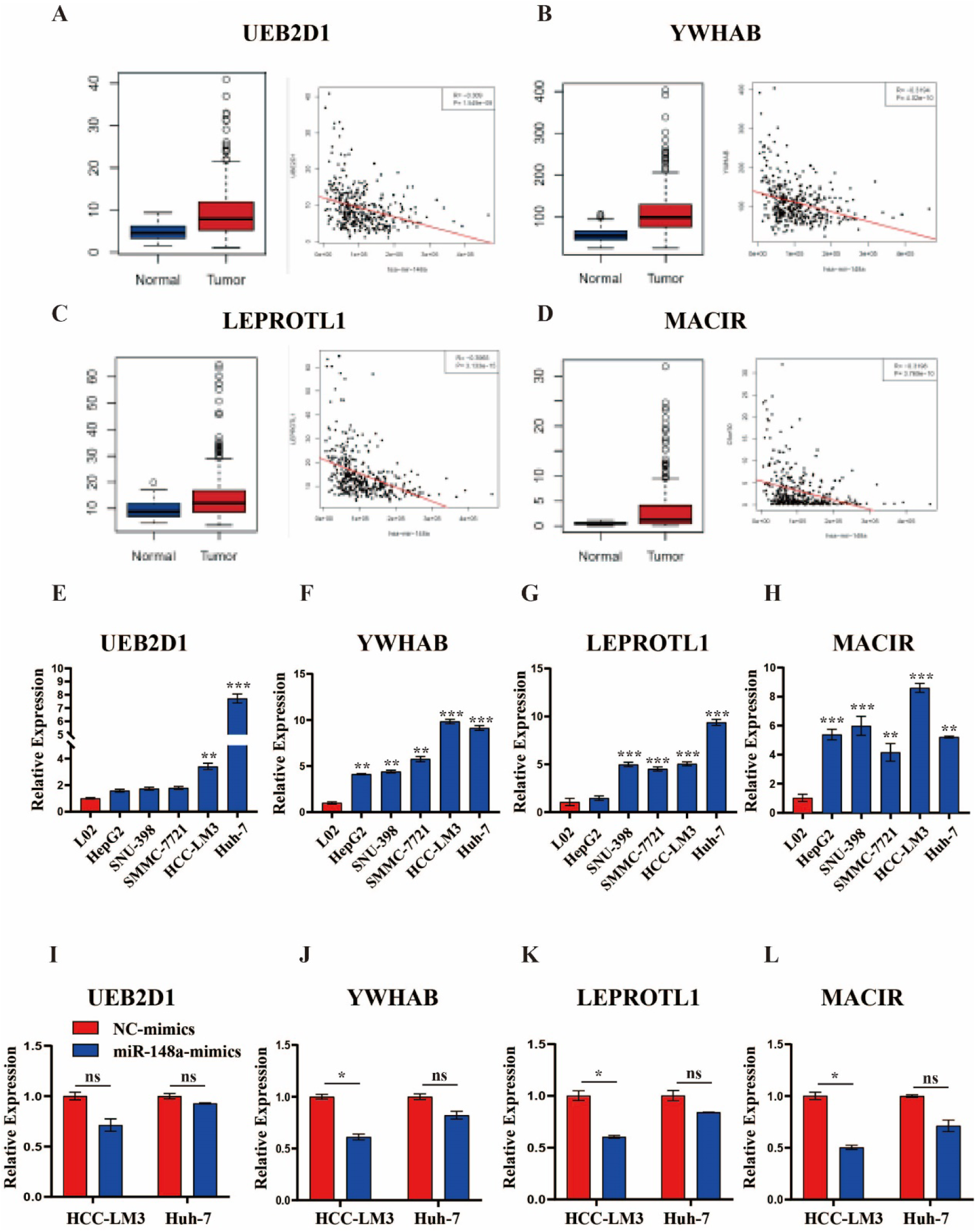
(A-D) TCGA analysis of UEB2D1, YWHAB, LEPROTL1 and MACIR expressions in HCC tissues and correlation analysis of these genes and miR-148a expressions. (E-H) RT-qPCR assays of UEB2D1, YWHAB, LEPROTL1 and MACIR expressions in HCC cell lines compared to L02 cells. (I-L) RT-qPCR assays of UEB2D1, YWHAB, LEPROTL1 and MACIR expressions in miR-148a overexpression HCC-LM3 and Huh-7 cells. Error bars represent the means ± SEM of 3 independent experiments. \**P* < 0.05, \*\**P* < 0.01, \*\*\**P* < 0.001.

**Figure S4.**
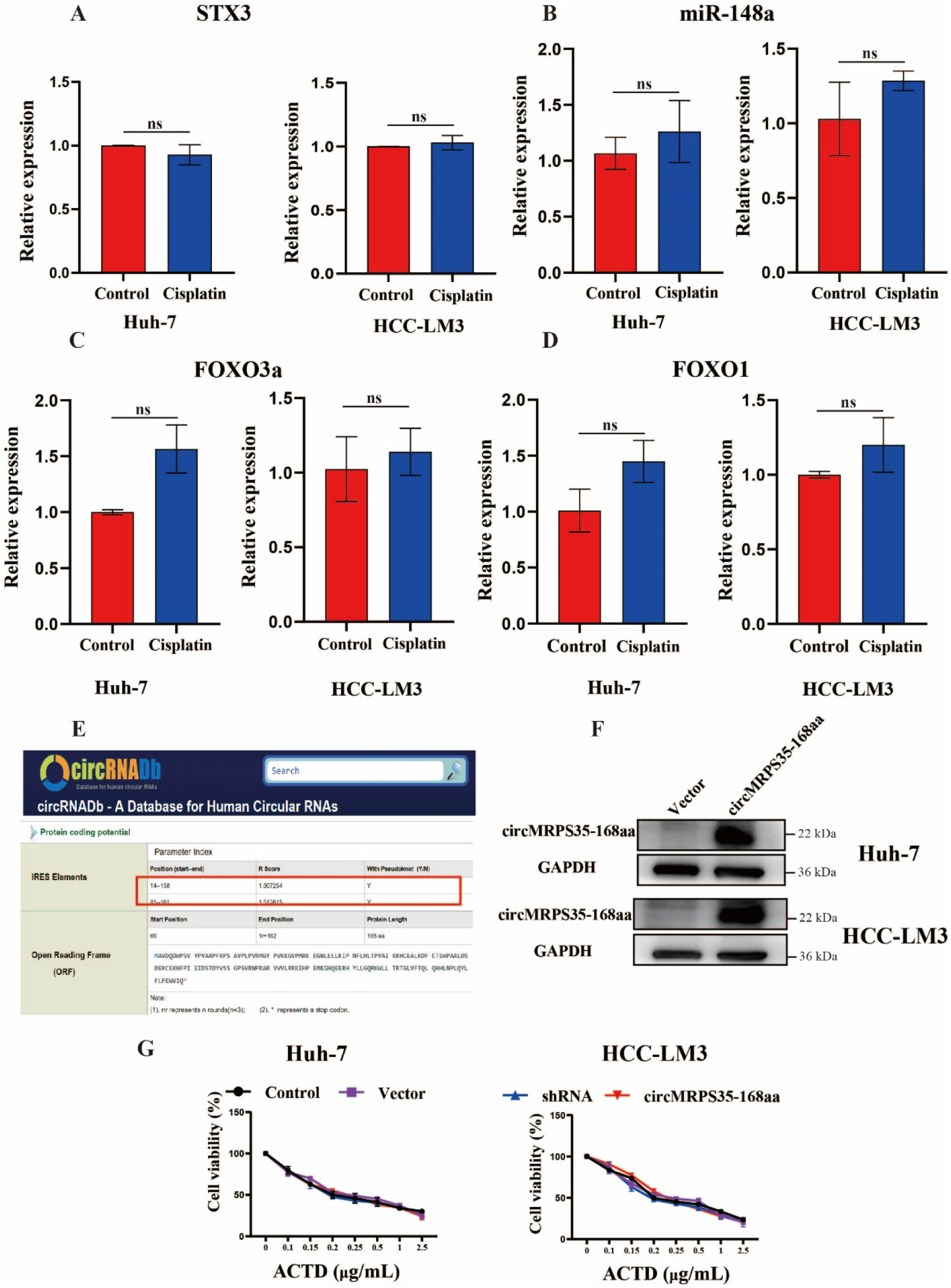
(A-D) RT-qPCR analysis of STX3, miR-148a, FOXO3a and FOXO1 expressions in cisplatin treatment or none-treated Huh-7 and HCC-LM3 cells. (E) circRNADb database showed IRES regions and the potentially peptide translated by circMRPS35. (F) Western blot analysis of circMRPS35-168aa in circMRPS35-168aa overexpression Huh-7 and HCC-LM3 cells, GAPDH as the positive control. (G) Cell viability assay of different circMRPS35-168aa expressed Huh-7 and HCC-LM3 cells with different concentrations of ACTD treatment. Error bars represent the means ± SEM of 3 independent experiments. \**P* < 0.05, \*\**P* < 0.01, \*\*\**P* < 0.001.

## Notes

### Competing Interest Statement

The authors have declared no competing interest.

## Reference

1. Bray, F, Ferlay, J, Soerjomataram, I, Siegel, RL, Torre, LA, and Jemal, A (2018). Global cancer statistics 2018: GLOBOCAN estimates of incidence and mortality worldwide for 36 cancers in 185 countries. CA Cancer J Clin 68: 394–424.

2. Gapstur, and M., S. Cancer Epidemiology and Prevention, 3rd Edition. Medicine & Science in Sports & Exercise 39: 395.

3. Clark, T, Maximin, S, Meier, J, Pokharel, S, and Bhargava, P (2015). Hepatocellular Carcinoma: Review of Epidemiology, Screening, Imaging Diagnosis, Response Assessment, and Treatment. Curr Probl Diagn Radiol 44: 479–486.

4. Mazzoccoli, G, Miele, L, Oben, J, Grieco, A, and Vinciguerra, M (2015). Biology, Epidemiology, Clinical Aspects of Hepatocellular Carcinoma and the Role of Sorafenib. Current Drug Targets 17.

5. Li, L, Chen, J, Chen, X, Tang, J, Guo, H, Wang, X, et al. (2016). Serum miRNAs as predictive and preventive biomarker for pre-clinical hepatocellular carcinoma. Cancer Lett 373: 234–240.

6. Xiao, Y, Liu, G, Sun, Y, Gao, Y, Ouyang, X, Chang, C, et al. (2020). Targeting the estrogen receptor alpha (ERalpha)-mediated circ-SMG1.72/miR-141-3p/Gelsolin signaling to better suppress the HCC cell invasion. Oncogene.

7. Wei, L, Wang, X, Lv, L, Liu, J, Xing, H, Song, Y, et al. (2019). The emerging role of microRNAs and long noncoding RNAs in drug resistance of hepatocellular carcinoma. Mol Cancer 18: 147.

8. Kanthaje, Shruthi, Makol, Ankita, Chakraborti, and Anuradha (2018). Sorafenib response in hepatocellular carcinoma: MicroRNAs as tuning forks. Hepatology Research the Official Journal of the Japan Society of Hepatology.

9. Houseley, JM, Zaida, GC, Maya, P, Nuria, P, O’Dell, KMC, Monckton, DG, et al. (2006). Noncanonical RNAs From Transcripts of the Drosophila muscleblind Gene. Journal of Heredity: 3.

10. Suzuki, H, and Tsukahara, T. A View of Pre-mRNA Splicing from RNase R Resistant RNAs. International Journal of Molecular Sciences 15: 9331–9342.

11. Salzman, J, Chen, RE, Olsen, MN, Wang, PL, Brown, PO, and Moran, JV. Cell-Type Specific Features of Circular RNA Expression. Plos Genetics 9: e1003777.

12. Han, D, Li, J, Wang, H, Su, X, Hou, J, Gu, Y, et al. (2017). Circular RNA circMTO1 acts as the sponge of microRNA-9 to suppress hepatocellular carcinoma progression. Hepatology 66: 1151–1164.

13. Zhu, YJ, Zheng, B, Luo, GJ, Ma, XK, Lu, XY, Lin, XM, et al. (2019). Circular RNAs negatively regulate cancer stem cells by physically binding FMRP against CCAR1 complex in hepatocellular carcinoma. Theranostics 9: 3526–3540.

14. Liang, WC, Wong, CW, Liang, PP, Shi, M, Cao, Y, Rao, ST, et al. (2019). Translation of the circular RNA circbeta-catenin promotes liver cancer cell growth through activation of the Wnt pathway. Genome Biol 20: 84.

15. Liu, Q, Cai, Y, Xiong, H, Deng, Y, and Dai, X (2019). CCRDB: a cancer circRNAs-related database and its application in hepatocellular carcinoma-related circRNAs. Database (Oxford) 2019.

16. Ding, Y, Fang, A, Yan, J, Duan, J, Wang, N, Yi, Y, et al. (2019). Selective downregulation of distinct circRNAs in the tissues and plasma of patients with primary hepatic carcinoma. Oncol Lett 18: 5255–5268.

17. Zhen, N, Gu, S, Ma, J, Zhu, J, Yin, M, Xu, M, et al. (2019). CircHMGCS1 Promotes Hepatoblastoma Cell Proliferation by Regulating the IGF Signaling Pathway and Glutaminolysis. Theranostics 9: 900–919.

18. Wang, B, Chen, H, Zhang, C, Yang, T, and Xu, F (2018). Effects of hsa_circRBM23 on Hepatocellular Carcinoma Cell Viability and Migration as Produced by Regulating miR-138 Expression. Cancer Biotherapy and Radiopharmaceuticals 33: 194–202.

19. He, J, Huang, Z, He, M, Liao, J, Zhang, Q, Wang, S, et al. (2020). Circular RNA MAPK4 (circ-MAPK4) inhibits cell apoptosis via MAPK signaling pathway by sponging miR-125a-3p in gliomas. Mol Cancer 19: 17.

20. Ha, M, and Kim, VN (2014). Regulation of microRNA biogenesis. Nature Reviews Molecular Cell Biology 15: 509–524.

21. Nan, H, Han, L, Ma, J, Yang, C, Su, R, and He, J (2018). STX3 represses the stability of the tumor suppressor PTEN to activate the PI3K-Akt-mTOR signaling and promotes the growth of breast cancer cells. Biochim Biophys Acta Mol Basis Dis 1864: 1684–1692.

22. Jie, M, Wu, Y, Gao, M, Li, X, Liu, C, Ouyang, Q, et al. (2020). CircMRPS35 suppresses gastric cancer progression via recruiting KAT7 to govern histone modification. Mol Cancer 19: 56.

23. Abe, N, Matsumoto, K, Nishihara, M, Nakano, Y, Shibata, A, Maruyama, H, et al. (2015). Rolling Circle Translation of Circular RNA in Living Human Cells. Sci Rep 5: 16435.

24. Yang, Y, Gao, X, Zhang, M, Yan, S, Sun, C, Xiao, F, et al. (2018). Novel Role of FBXW7 Circular RNA in Repressing Glioma Tumorigenesis. J Natl Cancer Inst 110.

25. Zhao, J, Lee, EE, Kim, J, Yang, R, Chamseddin, B, Ni, C, et al. (2019). Transforming activity of an oncoprotein-encoding circular RNA from human papillomavirus. Nat Commun 10: 2300.

26. Yang, Y, Fan, X, Mao, M, Song, X, Wu, P, Zhang, Y, et al. (2017). Extensive translation of circular RNAs driven by N(6)-methyladenosine. Cell Res 27: 626–641.

27. Cheng, Y, Sun, H, Wang, H, Jiang, W, Tang, W, Lu, C, et al. (2019). Star Circular RNAs In Human Cancer: Progress And Perspectives. Onco Targets Ther 12: 8249–8261.

28. Qiu, LP, Wu, YH, Yu, XF, Tang, Q, Chen, L, and Chen, KP (2018). The Emerging Role of Circular RNAs in Hepatocellular Carcinoma. J Cancer 9: 1548–1559.

29. Wang, M, Yu, F, and Li, P (2018). Circular RNAs: Characteristics, Function and Clinical Significance in Hepatocellular Carcinoma. Cancers (Basel) 10.

30. Yao, R, Zou, H, and Liao, W (2018). Prospect of Circular RNA in Hepatocellular Carcinoma: A Novel Potential Biomarker and Therapeutic Target. Front Oncol 8: 332.

31. Li, C, and Xu, X (2019). Biological functions and clinical applications of exosomal non-coding RNAs in hepatocellular carcinoma. Cell Mol Life Sci 76: 4203–4219.

32. Kristensen, LS, Andersen, MS, Stagsted, LVW, Ebbesen, KK, Hansen, TB, and Kjems, J (2019). The biogenesis, biology and characterization of circular RNAs. Nat Rev Genet 20: 675–691.

33. Meng, X, Li, X, Zhang, P, Wang, J, Zhou, Y, and Chen, M (2017). Circular RNA: an emerging key player in RNA world. Brief Bioinform 18: 547–557.

34. Hu, ZQ, Zhou, SL, Li, J, Zhou, ZJ, Wang, PC, Xin, HY, et al. (2019). Circular RNA Sequencing Identifies CircASAP1 as a Key Regulator in Hepatocellular Carcinoma Metastasis. Hepatology.

35. Terazawa, T, Kondo, S, Hosoi, H, Morizane, C, and Okusaka, T (2014). Transarterial infusion chemotherapy with cisplatin plus S-1 for hepatocellular carcinoma treatment: A phase I trial. Bmc Cancer 14: 301.

36. Ding, K, Fan, L, Chen, S, Wang, Y, Yu, H, Sun, Y, et al. (2015). Overexpression of osteopontin promotes resistance to cisplatin treatment in HCC. Oncol Rep 34: 3297–3303.

37. Ding, B, Lou, W, Xu, L, and Fan, W (2018). Non-coding RNA in drug resistance of hepatocellular carcinoma. Biosci Rep 38.

38. Luo, Y, Fu, Y, Huang, R, Gao, M, Liu, F, Gui, R, et al. (2019). CircRNA_101505 sensitizes hepatocellular carcinoma cells to cisplatin by sponging miR-103 and promotes oxidored-nitro domain-containing protein 1 expression. Cell Death Discov 5: 121.

39. Chen, H, Liu, S, Li, M, Huang, P, and Li, X (2019). circ_0003418 Inhibits Tumorigenesis And Cisplatin Chemoresistance Through Wnt/beta-Catenin Pathway In Hepatocellular Carcinoma. Onco Targets Ther 12: 9539–9549.

40. Guan, Y, Zhang, Y, Hao, L, and Nie, Z (2020). CircRNA_102272 Promotes Cisplatin-Resistance in Hepatocellular Carcinoma by Decreasing MiR-326 Targeting of RUNX2. Cancer Manag Res 12: 12527–12534.

41. Huang, XX, Zhang, Q, Hu, H, Jin, Y, Zeng, AL, Xia, YB, et al. (2020). A novel circular RNA circFN1 enhances cisplatin resistance in gastric cancer via sponging miR-182-5p. J Cell Biochem.

42. Zhang, J, Chen, S, Yang, J, and Zhao, F (2020). Accurate quantification of circular RNAs identifies extensive circular isoform switching events. Nat Commun 11: 90.

